# A subset of conserved phagocytic genes are likely used for the intracellular theft of cnidarian stinging organelles in nudibranch gastropods

**DOI:** 10.1101/2025.02.13.637864

**Authors:** Jessica A. Goodheart, Rose Fiorenza, Robin Rio, Rebecca N. Lopez-Anido, Noah J. Martin, Timothy J. Herrlinger, Rebecca D. Tarvin, Deirdre C. Lyons

**Affiliations:** Division of Invertebrate Zoology, American Museum of Natural History, New York, NY, USA; Scripps Institution of Oceanography, University of California, San Diego, La Jolla, CA, USA; Department of Integrative Biology, University of California, Berkeley, CA, USA; Museum of Vertebrate Zoology, University of California, Berkeley, CA, USA; Marine Biological Laboratory, Woods Hole, MA, USA

**Keywords:** nematocyst, sequestration, phagocytosis, immunity, defense, novelty

## Abstract

**Background:** Phagocytosis is a universal physiological process in eukaryotes with many important biological functions. In nudibranch gastropods, a novel form of phagocytosis called nematocyst sequestration is specialized for the uptake of venomous stinging organelles stolen from their cnidarian prey. This process is highly selective. Here we use the emerging model nudibranch species *Berghia stephanieae* and *Hermissenda opalescens* to identify genes enriched within the body regions where nematocyst sequestration occurs, and investigate how the expression profile of phagocytosis, immune, and digestive genes differs between nematocyst sequestering regions relative to those where other phagocytic functions occur.

**Results:** We identified 166 genes with significantly higher expression in sequestering regions in *B. stephanieae*, including genes associated with development, membrane transport, and metabolism. Of these, 41 overlap with transcripts upregulated in *H. opalescens* sequestering tissues. Using Hybridization Chain Reaction *in situs*, we show that at least two of these genes were localized to sequestering cells in *B. stephanieae*, including a putative C-type lectin receptor and a collagen. Genes annotated with phagocytosis, digestion, or immunity GO terms were often expressed in both sequestering and non-sequestering tissues, suggesting that they may also play a role in sequestration processes.

**Conclusion:** Our results suggest that phagocytosis genes likely play a role in the sequestration phenotype, and that a small subset of genes (e.g., collagen) may play unique functions yet to be uncovered. However, we also show that genes categorized in GO terms related to endocytosis, immunity, and digestion show a clear decrease in overall expression in sequestering tissues. This study lays the foundation for further inquiry into mechanisms of organelle sequestration in nudibranchs and other organisms.

## Background

Phagocytosis, or the process by which certain living cells ingest or engulf other cells or particles, is a universal physiological process across eukaryotes [1]. Phagocytosis is critical for basic biological functions including innate and adaptive immune responses, tissue homeostasis, and intracellular digestion [2,3]. Thus far, much of the work on phagocytosis has focused on “professional” phagocytes (i.e., cells where phagocytosis is the primary function, such as macrophages and neutrophils) from *in vitro* mammalian models and cell lines [4–6], along with a few other vertebrates [7] and ecdysozoans such as *Drosophila* [8] and *C. elegans* [9]. These studies largely center on immunity and homeostasis (anti-pathogen and clearance phagocytosis [10]). However, other phagocytes, such as digestive cells that perform intracellular digestion, are also common across Metazoa [11]. Among these groups, some nudibranch gastropods have evolved to steal stinging cells (i.e., nematocysts) from their cnidarian prey for defense, termed nematocyst sequestration [12,13].

In nudibranchs, nematocyst sequestration takes place at the distal edges of the digestive tract in an organ called the cnidosac, and within an evolutionarily novel cell type called a cnidophage [13–15]. This organ is at the distal end of the ceras (Fig. 1), which is made up of a number of tissue types, including neurosensory, epidermal, muscle, and digestive [18,21]. Some cell types are known to be found distributed across the ceras, including sensory cilial tufts and hemocytes (Fig. 1C). In the cnidosac, sequestered nematocysts appear to provide a defensive function [15–17] for the nudibranchs, as they do in their native cnidarians [18,19], but we know very little about how they are phagocytosed and stored. Previous research on phagocytosis in mollusks has focused on the bivalve immune response (e.g., within hemocyte cells [20,21]), including environmental effects on phagocytosis and immunity, but have rarely extended to molecular mechanisms [22–25]. Furthermore, mollusk immune cells are primarily generalists with regard to foreign targets, often expressing many different receptors in each cell [26]. In contrast, cnidophages exhibit a highly selective form of phagocytosis that exclusively targets nematocysts separated from their original cnidarian cnidocyte cell (Fig. 1C) [14].

**Figure 1.**
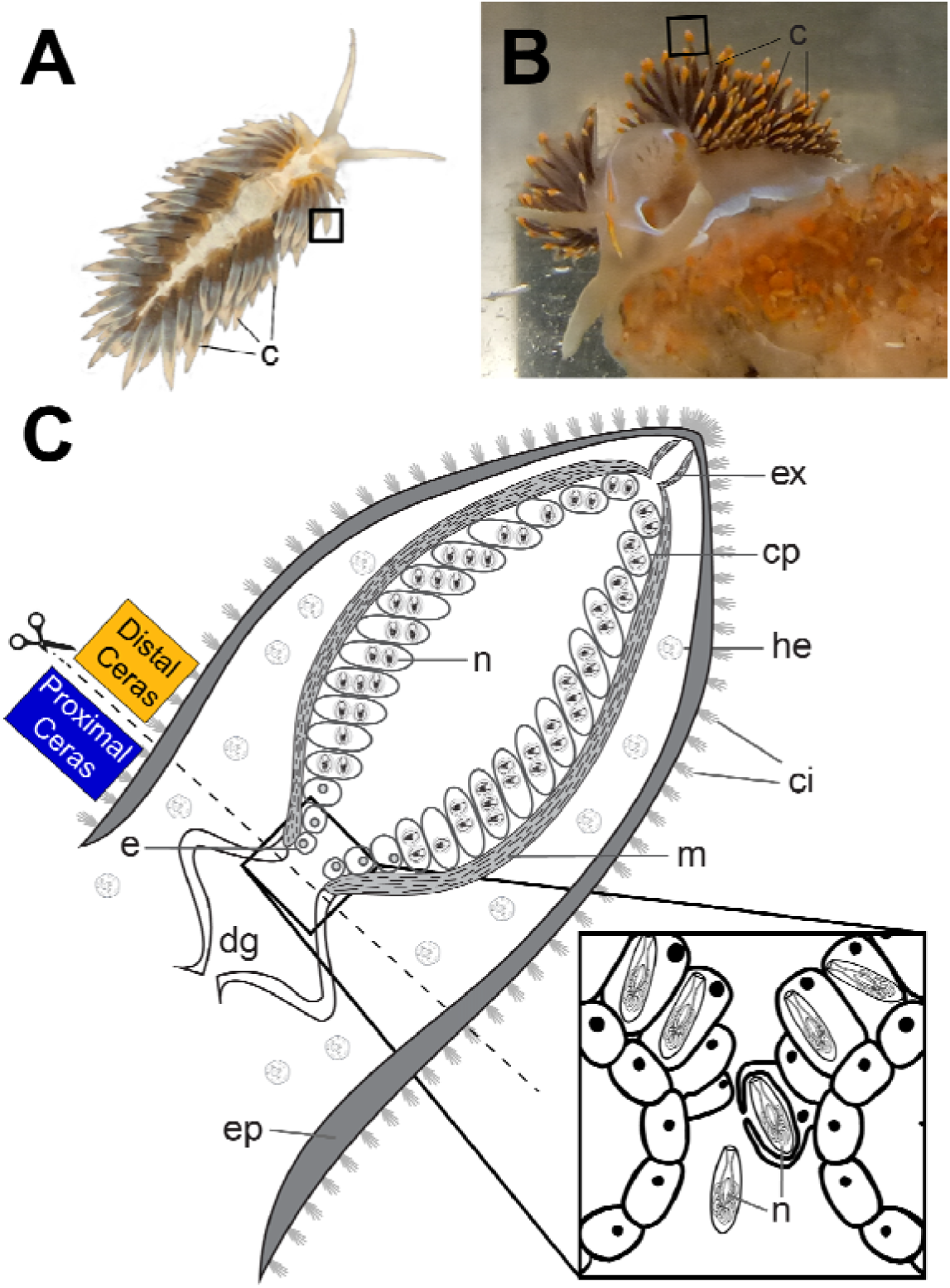
Specialized phagocytosis (i.e., nematocyst sequestration) of nematocysts in (A) *Berghia stephanieae,* and (B) *Hermissenda opalescens*. (C) Generalized cnidosac schematic modified from Goodheart *et al.* 2018 [14] (under <u>CC BY 4.0</u> Creative Commons License) highlighting the main features of the cnidosac. (*Inset)* Nematocysts (n) are phagocytosed by cnidophages inside the cnidosac (cs). Abbreviations: c, cerata; ci, cilia tufts; cp, cnidophage, dg, digestive gland; e, entrance to the cnidosac; ep, epithelium; ex, exit from the cnidosac; he, hemocytes; m, musculature; n, nematocysts.

Here we use the emerging model nudibranch species *Berghia stephanieae* (Valdés, 2005) [27] (Fig. 1A) and the Opalescent nudibranch *Hermissenda opalescens* (Cooper, 1863) (Fig. 1B) to identify candidate genes and processes used by cnidophages to steal stinging cells. We compare the expression of phagocytosis, immune, and digestive genes between nematocyst-sequestering regions (Distal Cerata) and non-sequestering digestive regions (Proximal Cerata) in adult and juvenile cerata tissues. The molecular machinery necessary for phagocytosis often includes a diverse array of recognition receptors, particle internalization proteins, and phagosome maturation proteins [28–30]. We would therefore expect some mechanistic conservation in the phagocytic processes of cnidophage cells compared to other digestive cell types. However, we do not expect all molecular signatures of traditional phagocytes to be present in the cnidosac, because cnidophages must identify nematocysts using a more specific form of target recognition [31], and phagosome maturation may be arrested to prevent nematocyst degradation. Instead, we hypothesize that we would find a narrowing of phagocytic, digestive, and immune function in cnidophages compared to other, more broad phagocytic tissues such as the digestive system, and that the cnidosac may express genes specific to targeting and engulfment of nematocysts. We predict that this will be in the form of phagocytosis, digestion, or immunity genes being downregulated in sequestering tissues.

## Results

### Differential expression across cerata tissues

The results obtained from each of the two species were collected under different experiments, so we present them separately here. For *B. stephanieae*, we used existing RNA-seq samples [32] for the distal and proximal cerata to perform our analyses (three of each), where the distal cerata were separated from proximal cerata just proximal to entrance of the cnidosac (Fig. 1C). The primary difference between the two regions is that the Distal Ceras contains the cnidosac, or the organ for nematocyst sequestration, and the Proximal Ceras contains primarily digestive tissues. The number of read pairs ranged from 23.7 million (SRX21326248, Distal Ceras) to 28.8 million (SRX8599772, Proximal Ceras) across all six samples (xlJ = 27.1 ± 2.3 million reads per sample; Table S1). On average, 75.9% (± 4.3%) of read pairs mapped uniquely to the *B. stephanieae* genome, and mapping percentage ranged from 68.5% of read pairs (SRX21326249, Proximal Ceras) to 80.6% of read pairs (SRX8599773, Distal Ceras). We identified expression (counts >10 across all tissues [33]) of ∼86.1% of genes (21,498 out of 24,960 predicted genes). Genes with a Log2 Fold Change > 2 and adjusted p-value < 0.05 were considered upregulated in a given tissue. Distal and Proximal Ceras tissues were found to have distinct expression profiles, though there were many similarities (Fig. S1-S2). We identified 166 genes with significantly higher expression in the Distal Ceras, including genes often associated with membrane structure and reorganization, such as Collagen, Actin, and alpha tubulin, genes associated with lysosome activity, such as Cathepsin L and Battenin, and genes associated with receptor-mediated endocytosis, such as C-type lectin Macrophage Mannose receptors and a Sortilin-related receptor (Table S3). By contrast, our analysis identified 458 genes with significantly lower expression in the Distal Ceras, including many putative phagocytosis genes such as C1q-like proteins and antigen-like proteins (Table S5). However, we also found extensive overlap in the genes expressed in the Distal Ceras and Proximal Ceras in *B. stephanieae* (∼87% of the total genes expressed, Fig. S3), including many other phagocytosis genes such as non-opsonic receptors, activation of internalization, phagosome formation and activation, phagocytosis-related responses, and phagocytosis efficiency genes (Table S2).

For *H. opalescens*, RNA-seq samples for the distal and proximal cerata ranged from 41.2 million (SRR32330747, Distal Ceras) to 96.5 million (SRR32330750, Distal Ceras) read pairs (xlJ = 27.1 ± 2.3 million reads per sample; Table S1). The constructed reference transcriptome for *H. opalescens* included 19,347 transcripts, with an average sequence length of 1,202 nucleotides and an N50 of 1,839. The metazoa_odb10 BUSCO analysis indicated 75.3% completeness (C:75.3%[S:71.4%,D:3.9%],F:10.6%,M:14.1%,n:954). On average, mapping rates were low (Table S1), with 42.1% (± 9.1%) of read pairs mapped uniquely to the existing *H. opalescens* transcriptome, and mapping percentage ranging from 26.8% of read pairs (SRR32330748, Proximal Ceras) to 58.7% of read pairs (SRR32330742, Distal Ceras). Due to these low mapping rates, and because *H. opalescens* data were more variable compared to our results in *B. stephanieae* (Fig. S3-S6), we used the *Hermissenda opalescens* results as a comparison to our findings in *B. stephanieae* rather than focusing our deep analysis efforts in this species. In *H. opalescens,* a total of 191 transcripts were found to be upregulated in the Distal Ceras, which were clustered within 159 orthologous groups by our OrthoFinder analysis. Thirty-six (36) of these orthologous groups (containing 71 *H. opalescens* transcripts and 41 *B. stephanieae* genes) were found to be upregulated in the Distal Ceras in both species (Fig. 2A, Table 1). The transcripts upregulated in the *H. opalescens* Distal Ceras include known phagocytosis genes, including those annotated as Collagen alpha-1(VIII) chain, Cathepsin L, multiple sodium-dependent transporter genes, and a C-type lectin macrophage mannose receptor.

**Figure 2.**
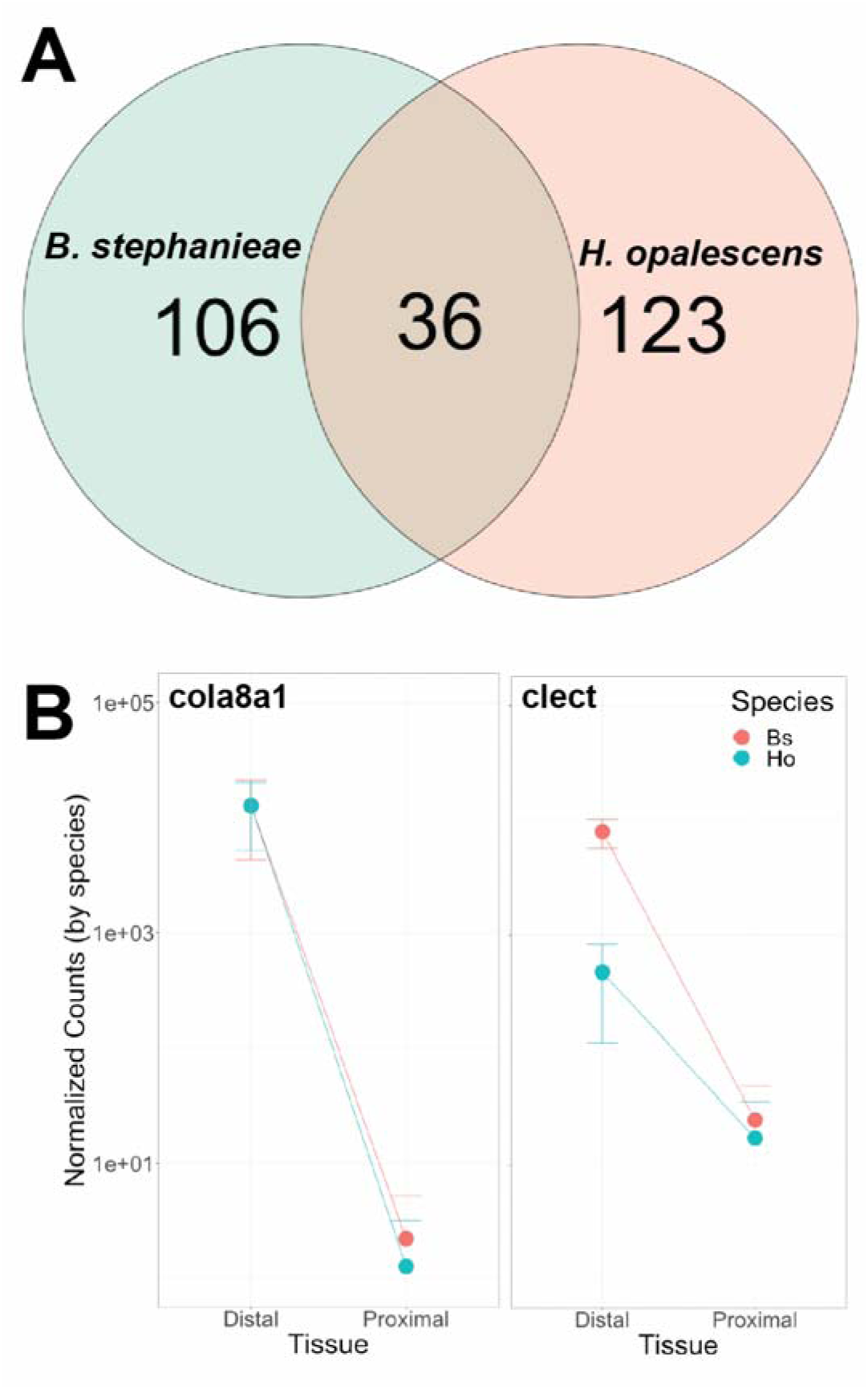
Select *Hermissenda* and *Berghia* gene expression comparisons. (A) Venn diagram indicating the proportion of orthologous groups containing genes upregulated in the Distal Ceras overlapped beteween species. (B) Plots showing gene expression levels between the distal and proximal ceras for two orthologous phagocytosis genes in *Berghia* and *Hermissenda*, including OG0034465 (Collagen alpha-1 chain, or cola8a1) and OG0000472 (Macrophage mannose receptor 1, or clect). Expression profiled for these two genes are shown in Figure 3.

**Table 1.**
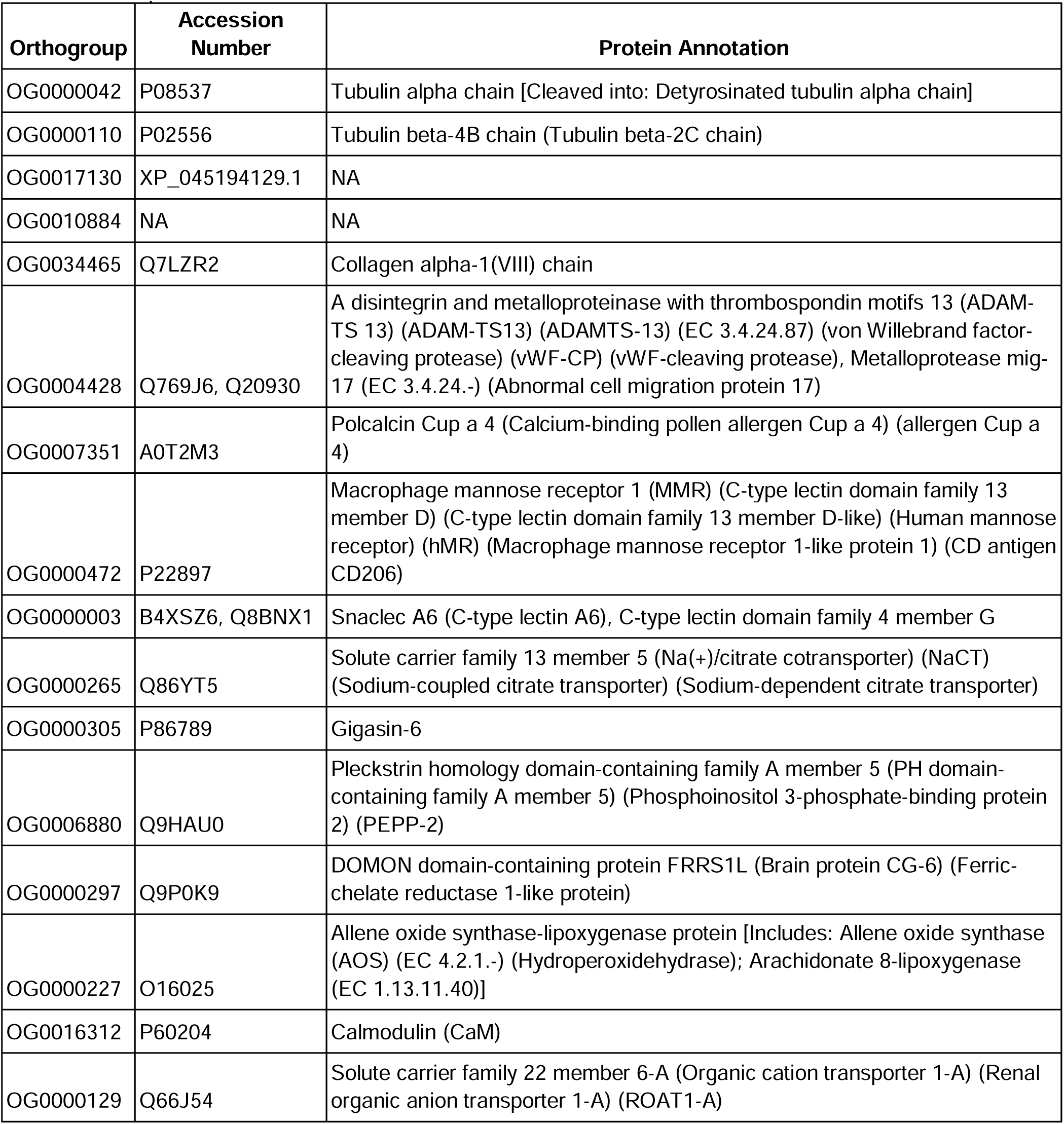
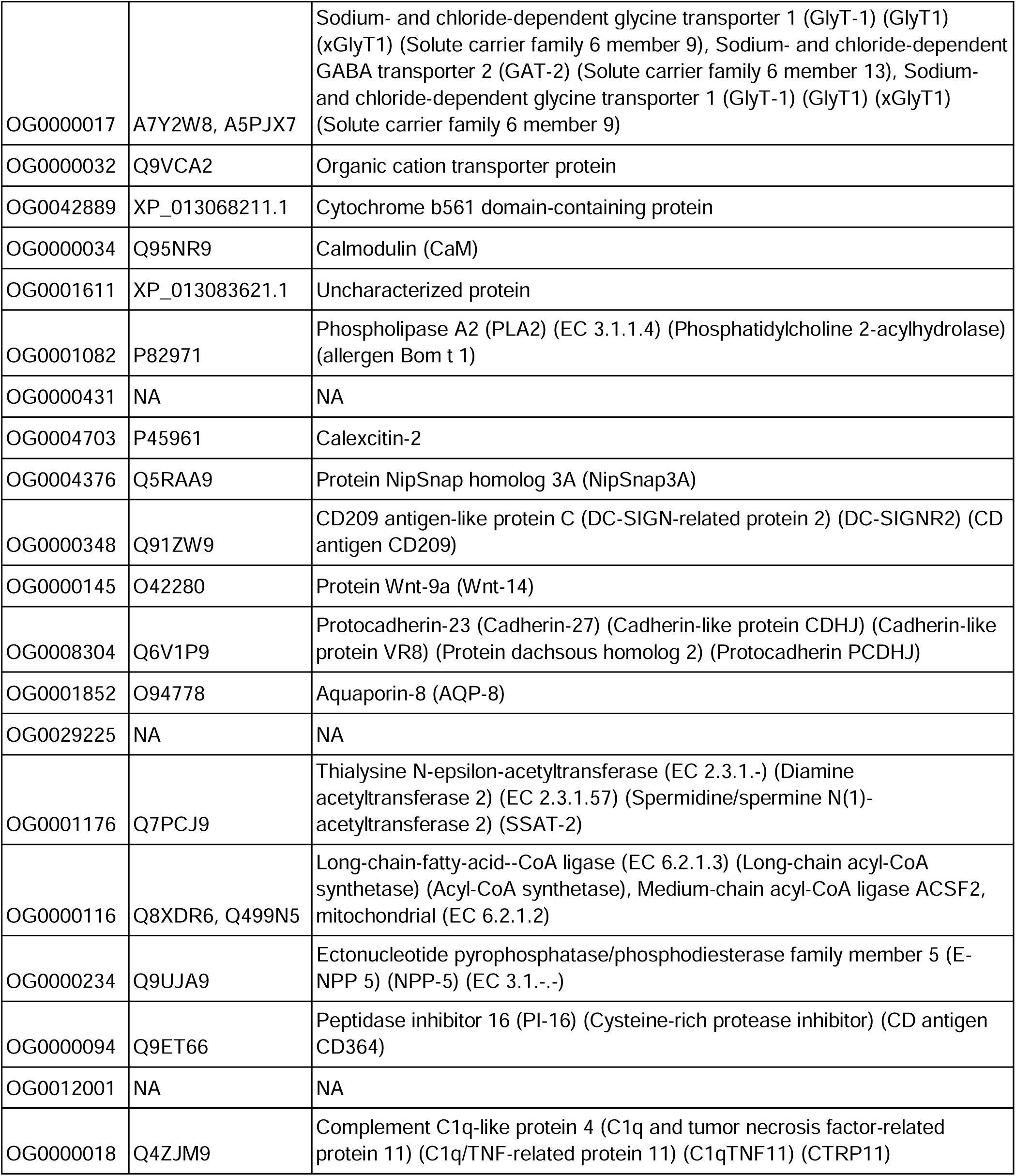
Shared upregulated genes in the Distal Cerata of *B. stephanieae* and *H. opalescens* clustered into orthologous groups. Also included are the Accession Number(s) of the best BLAST hits to NCBI and the associated protein annotation(s) from Goodheart *et al.* 2024 [32].

### Confirmed expression of upregulated genes in *B. stephanieae*

To further localize gene expression (Table 1), we used *in situ* Hybridization Chain Reaction (HCR) at different stages of cerata development in *B. stephanieae* [34]. First we examined the spatial expression of a known cnidophage marker in *B. stephanieae*, *bscola8a1* (OG0034465; annotated as collagen alpha-1 VIII), and that of a putative cnidophage marker *bsclect* (OG0000472; annotated as c-type lectin) in both juveniles and adults. The expression pattern of *bscola8a1* was described in juveniles previously [32]. Prior to the formation of cerata buds, we did not detect any expression of *bsclect* or *bscola8a1* (Fig. 3A-B). In later staged juvenile (2-3 days post-feeding) [31], we detected co-expression of *bsclect* and *bscola8a1* in the cnidosac at the onset of cerata development (Fig. 3C-D). Both *bsclect* and *bscola8a1* appear to have restricted expression in the cnidosac (Fig. 3C), with *bscola8a1* exhibiting a slightly stronger signal than *bsclect* (Fig. 3D). To detect expression at a finer cellular level in the adult *B. stephanieae*, we conducted HCR on paraffin-sectioned cerata tissue. We identified co-expression of *bsclect* and *bscola8a1* in cnidophages within the cnidosac of adult *B. stephanieae* (Fig. 3E-F), with *bscola8a1* again showing a stronger signal than *bsclect*. Together we found co-localization of *bsclect* and *bscola8a1* within the cnidosac at multiple stages of cerata development (Fig. 3C-F).

**Figure 3.**
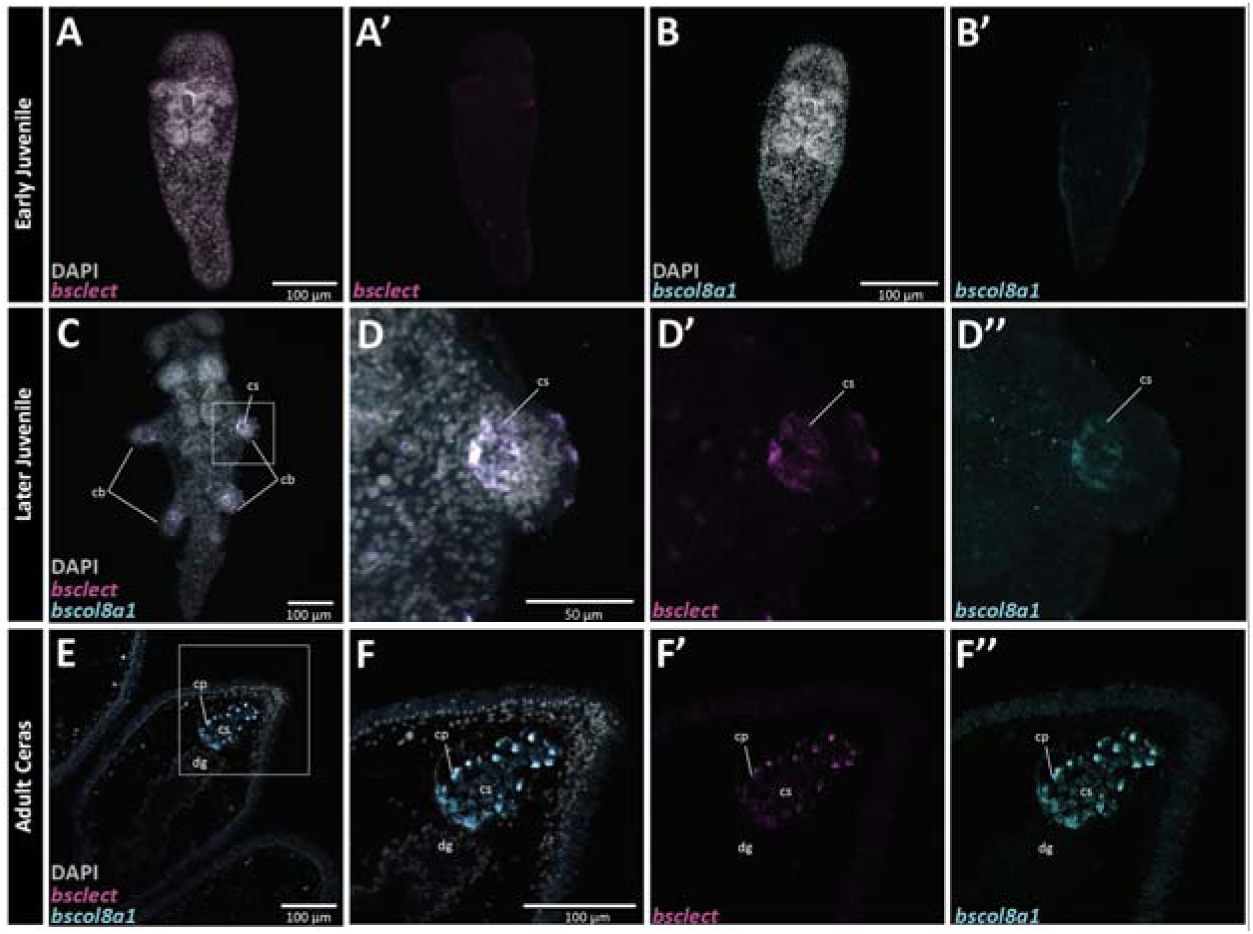
Hybridization chain reaction (HCR) detected expression of cnidosac markers (*bsclect* and *bscola8a1*) during cerata development in *Berghia stephanieae*. Early juvenile expression of (A-A’) *bsclect* and (B-B’) *bscola8a1* prior to cerata development. Later juvenile co-expression of (C-D’’’) *bsclect* and *bscola8a1* at the beginning of cerata development. Adult expression of (E-F’’’) *bscola8a1* and *bsclect* in the ceras tissue. Abbreviated: cb, cerata buds; cs, cnidosac; cp, cnidophage; dg, digestive gland.

To examine spatial expression of putative digestive gland genes, we selected *bsferritin* (jg36533; annotated as soma ferritin) and *bsbhmt* (jg32975; annotated as betaine--homocysteine S-methyltransferase 1) based on their upregulation in proximal cerata tissue from our differential expression analysis and their putative role in digestion [35] and metabolism [36], respectively. During early juvenile development (prior to the formation of cerata), we detected strong expression of *bsferritin* in the anterior portion of the digestive gland (Fig. 4A-B). In contrast, we detected very weak expression of *bsbhmt* in the posterior digestive gland tissue (Fig. 4C-D). Further along in juvenile development (2-3 days post-feeding) [31], we found expression of both *bsferritin* and *bsbhmt* throughout the entire juvenile digestive gland (Fig. 4E-H). We did not detect any expression of *bsferritin* nor *bsbhmt* in the distal cerata where the cnidosac is located (Fig. 4F’ and H’).

**Figure 4.**
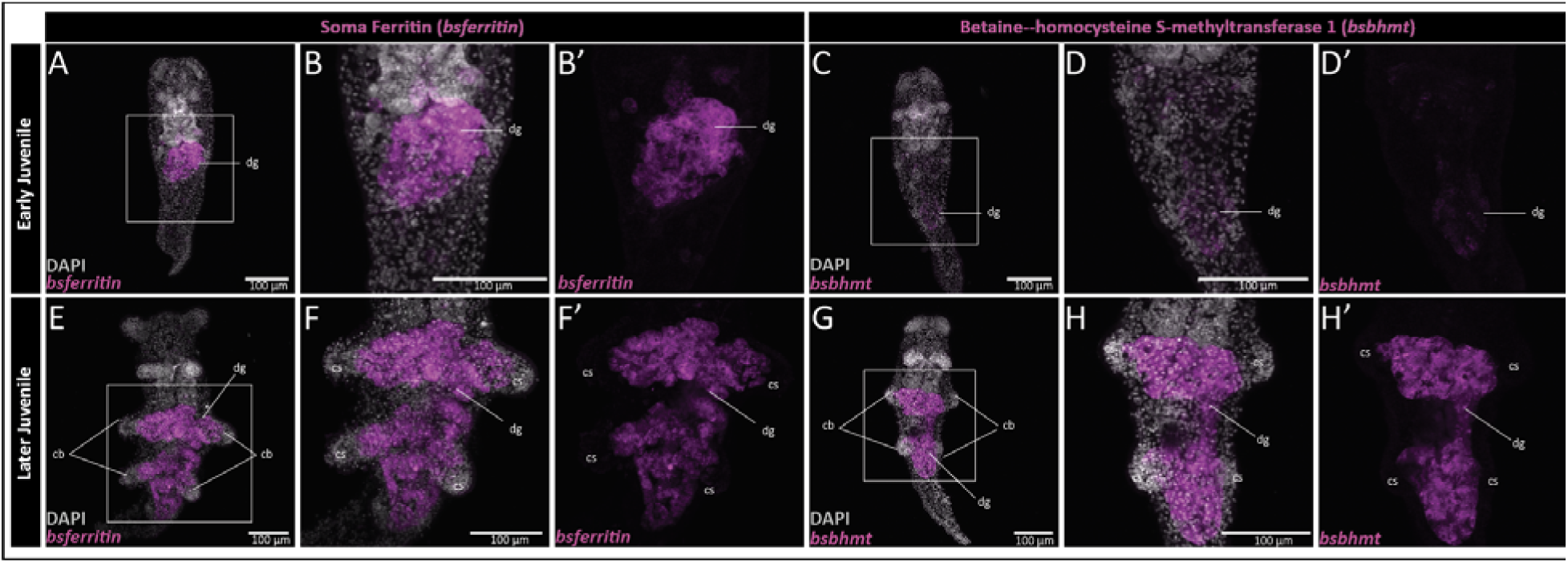
Hybridization chain reaction (HCR) detected expression of digestive markers (*bsferritin* and *bsbhmt*) during cerata development in *Berghia stephanieae*. Early juvenile expression of (A-B’) *BsFerritin* and (C-D’) *bsbhmt* prior to cerata development. Later juvenile expression of (E-F’) *bsferritin* and (G-H’) *bsbhmt* at the beginning of cerata development. Abbreviated: cb, cerata buds; cs, cnidosac; cp, cnidophage; dg, digestive gland.

To identify any background fluorescence and/or hairpin trapping, we conducted HCR with negative controls that were incubated without any HCR probes (Fig. S8). We found that both 546 and 647 hairpins get trapped in epidermal tissue, and this is particularly prominent in paraffin sectioned adult tissue (Fig. S8C-C’’).

### Predicted functional changes in the Distal Ceras

We predicted that a narrowing of function would occur in the Distal Ceras resulting in a downregulation of phagocytosis, digestion, or immunity genes in sequestering tissues. To test this hypothesis, we investigated the primary functions associated with GO terms of genes upregulated in the Distal Ceras compared to those downregulated in the Distal Ceras in *B. stephanieae.* Those upregulated in the Distal Ceras included some terms that make sense for cnidosac function, like transmembrane transport (Fig. 5A), but also included some very specific metabolic terms such as lactone metabolic process or cellular lipid metabolic process. The primary GO terms associated with genes downregulated in the Distal Ceras include those associated with immune response, endocytosis, and digestion (Table S6; Fig. 5B).

**Figure 5.**
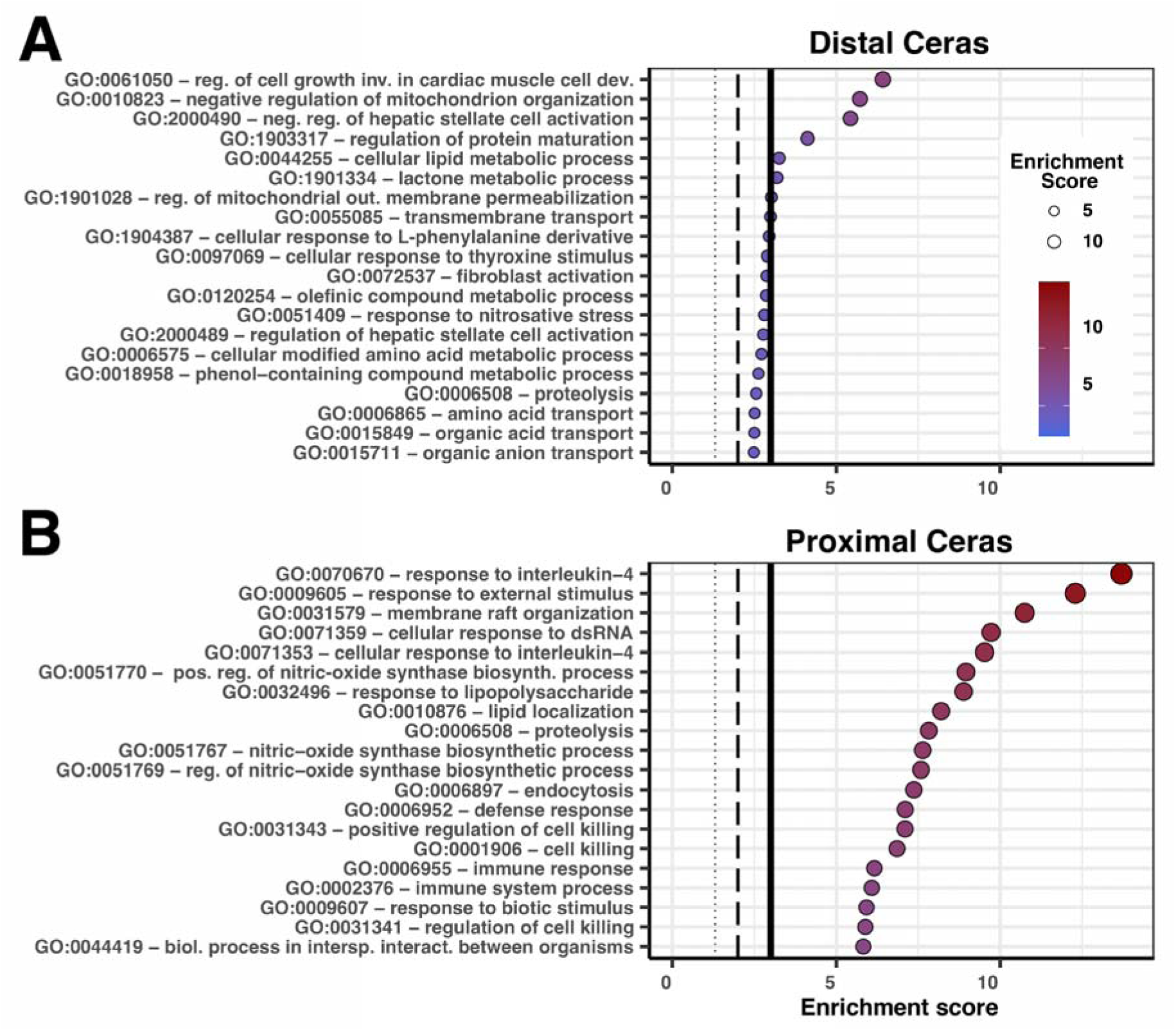
Plots describing the most common GO terms associated with genes upregulated in the (A) distal and (B) proximal cerata in B. stephanieae. We used the Fisher statistic with the ParentChild algorithm to determine which terms were significantly upregulated. The enrichment score is the −log10() of the calculated p-value. The lines in the plot represent the enrichment score that aligns with significance values at: p=0.05, small dotted line; p=0.01, middle dashed line; and p=0.001, thicker solid line. Enrichment scores are also represented in the size and color of the points, with larger and more red points representing lower p-values.

We further investigated genes related to Digestion (GO:0007586), Endocytosis (GO:0006897), and Immune Response (GO:0006955) GO terms by comparing normalized and transformed counts (see Methods for details) for all genes or only differentially expressed (DE) genes. We found that in these GO terms, the differentially expressed sets of genes were on average expressed at a lower level in the Distal versus Proximal ceras (Fig. 6). When comparing all orthologous genes across the Distal and Proximal Ceras, we found that these genes are not more likely to be up- or down-regulated in the Distal Ceras (Fig. 6A-C). However, we noted a clear pattern of directionality in differentially expressed genes, where these genes were more likely to be downregulated in the Distal Ceras compared to the Proximal Ceras (Fig. 6D-F).

**Figure 6.**
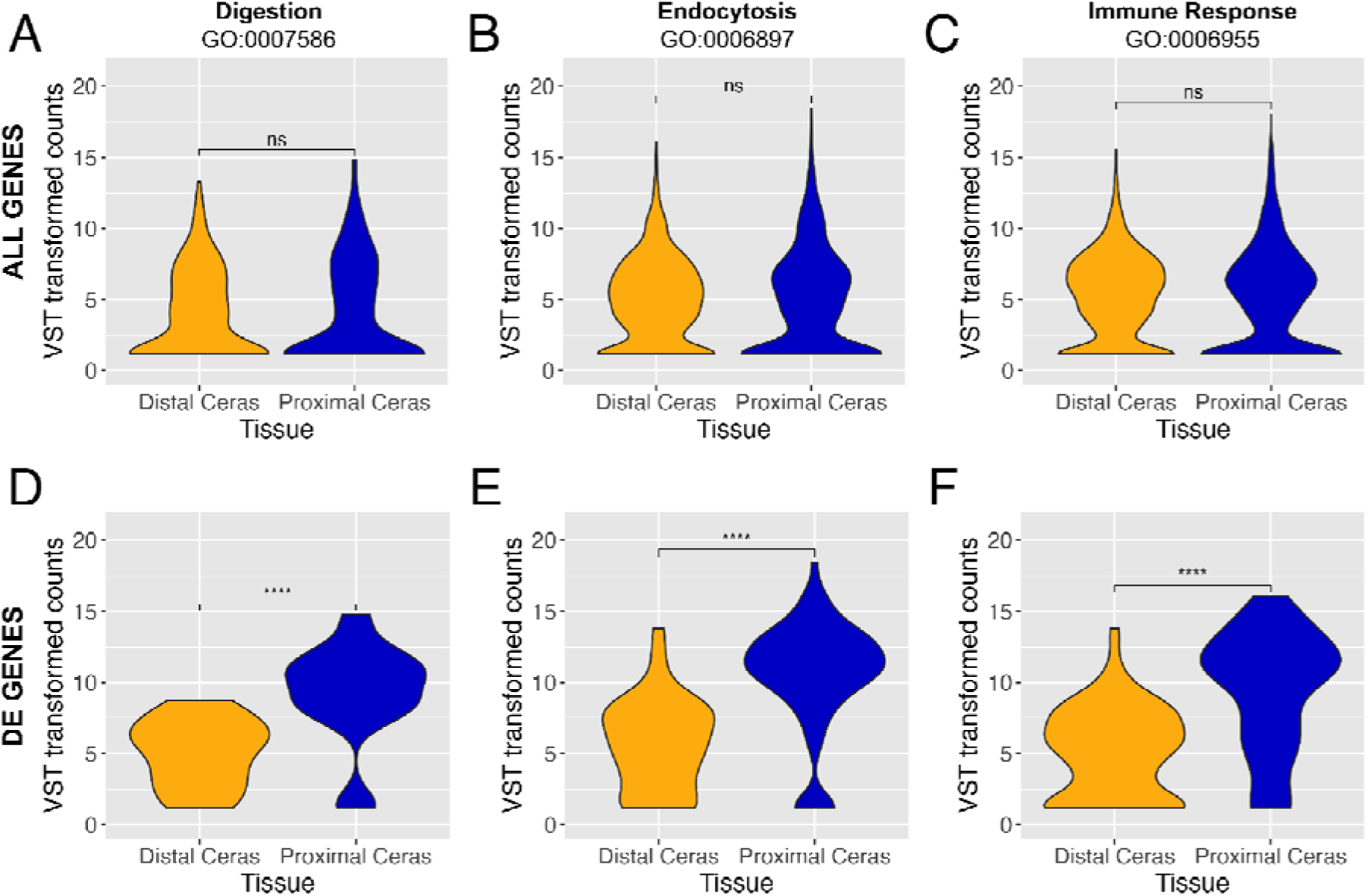
Violin plots comparing VST transformed counts of genes in our analysis assigned particular GO terms: (A,D) GO:0007586, Digestion (119 genes total; 9 DE genes); (B,E) GO:0006897, Endocytosis (525 genes; 33 DE genes); (C,F) GO:0006955, Immune Response (838 genes; 52 DE genes). Plots are for subsets of genes associated with each of these three GO terms: (A-C) for all genes associated with each GO term, and (D-F) for only differentially expressed (DE) genes across both tissues associated with each GO term. Asterisks indicate level of significance between the two tissues (ns, p > 0.05; *, p <= 0.05; **, p <= 0.01; ***, p <= 0.001; ****, p <= 0.0001).

## Discussion

Phagocytosis, a conserved process for ingesting large particles, has been described largely based on the function of immune cells in a few major model systems [37], and the extent to which these components are conserved or have diverged across the vast majority of metazoan diversity is not fully understood. Mollusk (and particularly gastropod) immune cells and phagocytes are especially useful for the study of phagocytosis, because many diseases, including schistosomiasis (caused by flatworms), can be indirectly transmitted to humans by these animals [38]. Mollusk models have also long been advantageous in studying cell types, disease, and basic cellular processes [39,40].

Here, we hypothesize a few crucial steps for nematocyst sequestration, a modified version of phagocytosis, to evolve in nudibranchs, including a narrowing of function identified by a reduction in expression of many genes related to phagocytosis, immunity, and common digestive functions. Prior research in *B. stephanieae* found that putatively “novel” genes are not more commonly expressed in distal cerata compared to other, more conserved tissues such as the nervous system [32]. This finding suggests that nematocyst sequestration is likely a mix of subfunctionalization and neofunctionalization [41], where a more generalist phagocytic cell type, likely digestive cells, evolved for specificity to nematocyst uptake and storage using largely conserved processes [31,32,42]. Our results provide further support for this hypothesis, showing that a body region (i.e., the Distal Ceras with the cnidosac) with a novel function (i.e., nematocyst sequestration) appears to express many genes associated with phagocytosis, digestion, and immunity, but with lower expression than is found in other body regions with more generalized phagocytic functions, like the Proximal Ceras (Fig. 5B, 6). We also show that nudibranch mollusks like *Berghia stephanieae* and *Hermissenda opalescens* offer opportunities for investigating how a conserved phenotype like phagocytosis results in a new phenotype like sequestration.

### Distal cerata tissues have lower expression of digestion and immunity genes

Cerata in nudibranchs are made up of a number of tissue types, including neurosensory, epidermal, muscle, and digestive [14,17]. Although the digestive gland and the cnidosac are only present in the proximal and distal cerata, respectively, many of the other functions of cerata are common to both tissue types. For example, cilial tufts are often distributed across the epidermis of the cerata, as are epidermal glandular cells [17], and hemocytes are often broadly distributed within the hemolymph (Fig. 1C). The functions of many of these glands remain unknown. It is unsurprising, therefore, that our results showed that the vast majority of genes expressed in the Proximal Ceras are also expressed in the Distal Ceras (87.3% overlap in *Berghia*). However, a clear separation in function exists between the proximal and distal cerata due to the presence of digestive gland tissue in the Proximal Ceras and the cnidosac in the Distal Ceras [14,31,43].

In the Proximal Ceras, the digestive gland is the primary internal tissue [14,44]. In the digestive gland, at least two epithelial phagocytic cell types have been described, typically regarded as digestive and basophilic cells [45]. Digestive cells are the most abundant cell type in the digestive gland, and are characterized by an endocytic activity and intracellular digestion [45]. Basophilic cells, by contrast, are considered the primary cell type for secretion of enzymes for extracellular digestion. Our results are consistent with these reported functions. For example, we see downregulation of genes associated with digestion and immune function in the Distal Ceras (Fig. 6; Table S6), with a higher expression of these genes in the Proximal Ceras compared to the Distal Ceras. We also identified some genes associated with digestion that are clearly only expressed in the digestive gland in *B. stephanieae*, including betaine—homocysteine S-methyltransferase (BHMT) and Soma Ferritin (Fig. 4), both of which are known to be used in mollusk digestion and metabolism [46,47]. The expression of many of the other genes more highly expressed in the Proximal Ceras still needs to be examined via *in-situ* hybridization methods to confirm exclusive expression in the digestive gland.

### Distal cerata tissues have higher expression of select conserved genes

We hypothesized that cnidophages, the primary cell type of nematocyst sequestration, would have a narrower phagocytic function compared to digestive cells, where fewer phagocytosis, digestion, and or metabolism genes are expressed. Our results support the hypothesis that nematocyst sequestration likely represents – in part – a subfunctionalization of digestive processes. This subfunctionalization has resulted in only a small subset of metabolic, immunity, and endocytic genes more highly expressed in the Distal Ceras compared to the Proximal Ceras in both *B. stephanieae* and *H. opalescens*, including genes such as Cathepsin L, Snaclec, alpha tubulin, and Calmodulin (Table 1). Included in this list are Collagen alpha-1(VIII) and Macrophage mannose receptor 1, which we found are expressed exclusively within the cnidosac in both species (Fig. 3).

Many phagocytosis mechanisms are still crucial to the function of nematocyst sequestration, including receptors that target nematocysts and genes involved with engulfment and phagosome formation. We hypothesized that we would identify a complement of receptors in the cnidosac that may be specific to nematocysts and thus relate to the neofunctionalization of the cnidosac. We found in *B. stephanieae* that a subset of these putative phagocytic receptors more highly expressed in the Distal Ceras compared to the Proximal Ceras, including multiple C-type lectin macrophage mannose receptors, Sortilin-related receptors, and cation-dependent mannose-6-phosphate receptors (Table S8). Our results with *B. stephanieae* also showed that Macrophage mannose receptor 1 (CLECT; Table S3, S8) was the putative receptor with the highest expression in the Distal Ceras, and we found that it is co-expressed with Collagen alpha-1(VIII) exclusively in the cnidophages (Fig. 3). Our interpretation is further supported by morphological work showing that digestive gland cells do not appear to endocytose and digest nematocysts in most nudibranchs, as nematocysts have only rarely been found within digestive cells in aeolid nudibranchs [18] and then only at the very distal parts of the digestive gland [58]. The lack of nematocyst endocytosis in the digestive gland may be because nematocysts must be able to reach the cnidosac without being destroyed by digestive cells, and suggests that some level of neofunctionalization has to have occurred in the cnidosac. However, previous work in Hancockiidae nudibranchs, a group representing a second origin of nematocyst sequestration in nudibranchs, has shown that nematocysts are endocytosed within the digestive gland in addition to the cnidosac. This difference between aeolids and dendronotid nudibranchs [42] provides an important distinction that would allow us to further test this hypothesis.

Outside of their need for target specificity, cnidophages still need to perform other conserved phagocytosis functions, such as the engulfment of the material and phagosome formation, which requires the conserved actin-myosin contractile system [48] and the microtubule system [49]. We found alpha-tubulin upregulated in both *B. stephanieae* and *H. opalescens* (Table 1) and both actin and myosin upregulated in the Distal Ceras in *B. stephanieae* (Table S3), which is in agreement with our assertion. However, it is possible that some of these genes produce different protein isoforms that have different functions between the two tissues [50], which would not be detectable in our analysis. Finally, we cannot yet reject the hypothesis that cnidophages are not the source of this difference; further analysis comparing expression profiles of cnidophage cells with that of other phagocytes (using techniques such as single-cell RNA-seq) are still necessary.

### The role of conserved genes in nematocyst sequestration

The role of conserved genes in nematocyst sequestration is still not fully resolved, but our data suggest that conserved genes are likely critical to the novel function of nematocyst sequestration in cnidophages. In other systems where cells or structures are engulfed and stored intracellularly, such as cnidarian-dinoflagellate endosymbiosis, it is clear that cnidarian cells use pattern recognition receptors (PRRs) to identify microbe-associated molecular patterns like peptidoglycans and lipopolysaccharides [51]. Mollusks have also undergone a large-scale expansion and molecular divergence of PRRs and signaling adapters, as well as significant recombination of immune receptors and fibrinogen-related proteins FREPs [52]. This may be part of the reason that mollusks like nudibranchs have evolved multiple types of phagocytic specializations, including nematocyst sequestration [43], kleptoplasty [53], and endosymbioses [54,55]. We found PRRs, and particularly Mannose receptors, expressed in the Distal Ceras of both nudibranch species (Table S8), and expressed exclusively within the cnidophages of *B. stephanieae* (Fig. 5), which supports this hypothesis.

However, some of these conserved genes may not necessarily retain precisely the same functions across phagocytic tissues and across large phylogenetic distances. For example, evidence suggests that in cnidarian-dinoflagellate endosymbiosis, activation of the host Transforming growth factor beta (TGFβ) innate immune pathway actually promotes tolerance of the symbiont rather than destruction [56]. This is consistent with the function of TGFβ immune regulation in mammals, as this pathway plays an important role in maintaining peripheral tolerance against self- and innocuous antigens [57]. We, and others [58], have found no evidence to support the role of this pathway in maintenance of sequestered nematocysts or symbionts in *B. stephanieae* or *H. opalescens*. It is clear that conserved genes are important in the evolution of novelties like sequestration and endosymbiosis, but the mechanisms driving the recruitment and co-option of certain genes and pathways are complex, and may be different in systems where specific organelles are the primary targets.

## Conclusions

Nematocyst sequestration, a process by which stinging organelles from cnidarians are stolen and stored intracellularly, is a novel function that relies on highly conserved phagocytic functions. As such, it is an excellent system in which to investigate how conserved functions can be modified to produce novel phenotypes. Our results clearly indicate that Distal Ceras tissues have very high expression of certain genes known to be associated with phagocytosis, including at least one that may be used for targeting nematocysts that we found expressed in the cnidophage cells. We also found that sequestering tissues tend to have lower expression of genes associated with digestion compared to digestive tissues. Although the role of conserved genes in nematocyst sequestration is still not fully resolved, our data suggests that many are critical to nematocyst sequestration despite the novel nature of this physiological process. From these results, we hypothesize a more detailed model for the evolution of the molecular mechanism of nematocyst sequestration, including: (1) the evolution of a means of specificity for targeting nematocysts over other tissues and structures, (2) modifications to internalization and phagosome formation pathways to accommodate multiple large and unusually shaped objects, and (3) the loss of some functions related to phagosome maturation, immunity, and common metabolic functions. This study lays the foundation for further inquiry into mechanisms of organelle sequestration in nudibranchs and other organisms.

## Methods

### Specimen collection

*Hermissenda opalescens* were collected from the wild at the Monterey Municipal Wharf #2 (36.60398, −121.88898) on 4 April 2019 and 14 September 2019 under permit SC-3726 from the State of California Department of Fish and Wildlife to TJH.

We maintained adults of *Berghia stephanieae* in continuous culture in the Lyons Lab at the Scripps Institution of Oceanography (SIO) or the Goodheart Lab at the American Museum of Natural History (AMNH), broadly following techniques laid out by Goodheart et al. 2022 and 2024 [31,32]. The original *Berghia stephanieae* animals obtained for culture were purchased from the company ReefTown. At SIO, we purchased *E. diaphana* from Carolina Biological Supply, Burlington, NC; at AMNH we purchased *E. diaphana* from the Marine Biological Laboratory, Woods Hole, MA.

### Transcriptomic data analysis

We obtained *Berghia stephanieae* Distal (3 samples; SRR12072207, SRR25598596-SRR25598597) and Proximal Ceras (3 samples; SRR12072208, SRR25598594-SRR25598595) read mapping data from the previously published *B. stephanieae* genome paper [32] through NCBI. To collect Proximal and Distal Ceras samples for *B. stephanieae*, we removed cerata and separated the region with the cnidosac (Distal Ceras) from the region with the digestive gland (Proximal Ceras) at the narrow entrance to the cnidosac (Fig. 1C) [14]. Transcriptome data for *H. opalescens* (SRR32330727-SRR32330757) was obtained from an experiment in which cerata were removed from field-collected animals and allowed to regrow. During the regrowth period, nudibranchs were separated into two experimental groups and maintained at 14°C in separate chambers. One group was fed daily with *Aurelia aurita* (moon jelly) polyps; the other was fed a non-nematocyst food source such as mussel, squid, or tunicate. Once the cerata were regrown (30-40 days later, [59]), we removed all cerata from the right side of the body and sectioned each into three parts: (1) the cnidosac (“Distal Ceras”), (2) “cer,” the region from between the cnidosac to about ⅔ the distance towards the body of the animal, and (3) “sub” (“Proximal Ceras”), the remaining region closest to the body of the animal. At this time, we also sampled foot tissue (i.e., not within the cerata) from each nudibranch. Total RNA was extracted from 5-10 pooled cerata tissues (separated by individual and section type) using TRIzol (Life Technologies), and mRNA was reverse transcribed using NEXTflexTM Poly(A) Beads (Bioo Scientific) and SuperScript® III Reverse Transcriptase (Life Technologies). The resulting cDNA was made into libraries using the NEXTflex Directional RNAseq Kit and sent for two lanes of 2x150 paired-end sequencing on the Illumina HiSeq4000 at QB3 Genomics (UC Berkeley, Berkeley, CA, RRID:SCR_022170).

For reference transcriptomes, we used the unfiltered BRAKER2 gene prediction (59,494 genes, 61,662 proteins) in the published *Berghia stephanieae* genome paper [32] as our *B. stephanieae* reference, and downloaded *Hermissenda opalescens* transcriptome data from NCBI (SRR1950939, [60]) to build our *H. opalescens* reference transcriptome. The *H. opalescens* reference transcriptome was generated using the protocol laid out in Goodheart et al. (2024) [32]. Briefly, using the previously assembled version of the *H. opalescens* transcriptome [61], we predicted open reading frames (ORFs) with TransDecoder (version 5.5.0; [112]) and clustered predicted ORFs using CD-HIT-EST (version 4.8.1; [113, 114]) at 95% identity and word size of 11 (-c 0.95, -n 11). Post-clustering, we filtered transcripts with alien_index (https://github.com/josephryan/alien_index); based on an algorithm described in [115]). We removed all sequences with an alien index greater than 45 from the transcriptome. We then compiled full transcripts for each predicted ORF sequence remaining from the assembled transcriptome using a custom Python script (full_transcripts.py, [32]).

We mapped reads to the *B. stephanieae* genome using STAR v2.7.9a [62] with default parameters plus additional flags (--readFilesCommand zcat --outSAMtype BAM SortedByCoordinate --twopassMode Basic --sjdbGTFfeatureExon ‘CDS’). We mapped reads to the *H. opalescens* reference transcriptome using Bowtie 2 v.2.4.4 [63]. We counted reads using the command htseq-count from the HTSeq framework v1.99.2 [64]. We analyzed counts using the DESeq function from DESeq2 v1.26.0 [65] to perform differential expression analysis, and generated the results using the results function. DESeq2 uses the median of ratios approach to normalize the counts data taking into account sequencing depth and RNA composition. We considered genes upregulated if the adjusted p-value (padj) was greater than 0.05 and log2FoldChange was greater than 2. Genes were considered exclusively expressed in a certain tissue if the average number of reads across all samples was >0.5 in that tissue and <0.5 in the other, which would mean that gene had a count of 1 in at least two samples. Orthology of *B. stephanieae* genes to *H. opalescens* transcripts was assigned using output from the OrthoFinder [66,67] analysis in Goodheart et al. 2024 [32].

Putative gene function was assigned based on GO terms. To assess which GO terms were significantly enriched in our two tissues, we used the Fisher’s Exact test implemented in the R package TopGO [68]. To directly compare expression of genes that fall under particular GO terms (Table S7), we performed a paired t-test implemented in ggpubr [69] that compared variance-stabilizing transformation (VST) counts (transformation performed on DESeq2 normalized counts) from the same gene between Proximal and Distal Cerata. VST transformation counts were used to make the data homoskedastic, and VST was selected because the range of size factors was < 4 [65]. These comparisons were used to evaluate whether genes associated with certain GO terms were more likely to be highly expressed in one set of tissues over another, particularly those that were considered differentially expressed.

### Tissue Fixation and Paraffin Section Preparation

We cultured and relaxed *B. stephanieae* juveniles using the same methods from prior *B. stephanieae* imaging work [31,32,70]. Prior to fixation, we starved juveniles for 3 days. Post-relaxation, we fixed juveniles using 4% Paraformaldehyde (PFA, diluted in Filtered Sea Water from 16% ampules). We washed samples in a 50% 1X PBS / 50% Methanol solution for 10 minutes prior to three washes in 100% Methanol for 10 minutes, all at room temperature. We stored samples in 100% Methanol at −20C for up to three months prior to HCR experiments. For adult *B. stephanieae*, we incubated individuals at 4C until fully relaxed and then fixed adults with a 4% PFA (diluted in DEPC-treated 1XPBS) incubation overnight at 4C. We washed and stored samples following the same protocol described for juveniles. Fixed samples that were put into paraffin immediately were dehydrated into 75% ethanol instead of 100% methanol.

To prepare fixed adult *B. stephanieae* for paraffin wax embedding (paraplast wax), we first placed the samples into 100% ethanol with two 100% ethanol washes for 2 minutes. We cleared the samples with three 20 minute washes in Histosol on a shaker at room temperature. We then washed samples in 50% Histosol 50% paraffin wax twice for 30 minutes each at 60 C. We left the samples in 100% paraffin wax overnight at 60 C, and then performed four 1 hour paraffin washes at 60 C before embedding the sample in paraffin using a mold. We sectioned samples using a Leica Biosystems RM2245 microtome to a thickness of 10uM. We floated sectioned samples in water before mounting onto Fisherbrand™ Superfrost™ Plus slides (FisherScientific #22-037-246). We then incubated the samples at 37 C overnight before storing them at room temperature until further use.

### Hybridization Chain Reaction

We designed all HCR probe sets using the HCR 3.0 probe maker with either the B1 or B2 amplifier (ÖzpolatLab-HCR, 2021, Kuehn et al., 2021) for up to 30 probe pairs (Tables S9-S13). We ordered probe sets (50LJpmol DNA oPools Oligo Pool) from Integrated DNA Technologies (Coralville, IA), which we resuspended to 1LJpmol/μl in 50LJmL TE buffer or RNase-free ultrapure water and rna and stored at −20C.

For whole-mount juvenile HCR, we used a modified HCR 3.0 protocol for *Berghia stephanieae* using the same solution recipes from Choi et al. (2018) [34] and modifications according to Goodheart et al. (2024)[32]. Following the hoechst staining, we mounted samples in a 20% 5X SSCT, 80% Glycerol solution and we followed the imaging process described in Goodheart et al. (2024) [32]. For HCR on paraffin sections, we followed the protocol from Choi et al. (2018) [34] with the modifications described in Criswell & Gillis (2020) [71]. We used the same probe and amplifier concentrations described in Goodheart et al. (2024) [32] for both whole-mount and paraffin section HCR.

### Fluorescent Imaging

We imaged whole-mount HCR samples with a Zeiss LSM 710 inverted confocal microscope with a AxioCamHRm camera. We imaged paraffin section HCR samples with a Zeiss LSM.

We analyzed and processed images with software ImageJ FIJI and AdobePhotoshop [72] (Adobe Inc., California, USA). We stitched images together using the FIJI Pairwise Stitching Plugin [73]. We created figures in Adobe Illustrator (Adobe Inc., California, USA).

## Supporting information

Tabls S2

Tabls S3

Tabls S4

Tabls S5

Tabls S6

Tabls S7

Tabls S8

Tabls S9

Tabls S10

Tabls S11

Tabls S12

Tabls S13

## List of Abbreviations

HCR: Hybridization Chain Reaction

## Declarations

### Ethics approval and consent to participate

Not applicable.

### Consent for publication

Not applicable.

### Competing interests

The authors declare that they have no competing interests.

### Availability of data and materials

Raw sequencing data used are accessible through the NCBI Sequence Read Archive (BioProjects PRJNA1004233, PRJNA641185, and PRJNA1222994), including for the Distal (3 samples; SRR12072207, SRR25598596-SRR25598597) and Proximal Ceras (3 samples; SRR12072208, SRR25598594-SRR25598595) samples for *Berghia stephanieae* and samples from multiple tissue types in *Hermissenda opalescens*. The genome used is available at DDBJ/ENA/GenBank under the accession JAWQJI000000000. Input and intermediate files for our analyses are available in Dryad (https://doi.org/10.5061/dryad.8pk0p2p00) and custom R scripts used in our analyses are available through Github (https://github.com/goodgodric28/sequestration_genes/tree/main/sequestration_genes).

### Funding

This work was supported by a Scripps Postdoctoral Fellowship and startup from the AMNH to JAG, and NIH BRAIN awards U01-NS108637 and 1U01-NS123972, an Emerging Research Organisms Grant from the Society for Developmental Biology to DCL, start-up funding from UC Berkeley to RDT, and a 2019 Conchologists of America Academic Grant to NM. RDT is also supported by NIH NIGMS R35GM150574. This work also used the Extreme Science and Engineering Discovery Environment (XSEDE) at the San Diego Supercomputer Center (SDSC) through allocation IDs TG-BIO210019 and TG-BIO210138 [74], which is supported by National Science Foundation grant number ACI-1548562, as well as the Savio computational cluster resource provided by the Berkeley Research Computing program at the University of California, Berkeley (supported by the UC Berkeley Chancellor, Vice Chancellor for Research, and Chief Information Officer). Sequencing at QB3 Genomics on the Illumina HiSeq 4000 platform was supported by NIH S10 OD018174 Instrumentation Grant.

### Authors’ contributions

JAG, NM, TDT, and DCL conceived of the study; JAG, RAF, RDT, NM, TJH, and RLA collected data; JAG, RDT, and RAR performed data analyses; all authors contributed to study design, participated in data interpretation, and read and approved the final manuscript.

## Acknowledgements

We thank all members of the Goodheart, Lyons, and Katz labs for help with animal care and feedback on this project and manuscript, particularly M. Desmond Ramirez, who was instrumental in helping us establish HCR in juveniles. We thank the Holford and Gillis labs for helping us establish paraffin section HCR in *B. stephanieae*. We are also grateful to members of the Berghia Brain Project (https://sites.google.com/umass.edu/berghiabrainproject/) for their feedback on this project. We also thank the Hamdoun Lab and the AMNH Microscopy Imaging Facility for the use of their confocal microscopes, the Rouse Lab for the use of their Qubit, and the AMNH Institute for Comparative Genomics for assistance with the laboratory portion of this work. We also thank Connor Tumelty, Yin Chen Wan, Kristen Tamsil, and Kate Montana for assistance with *H. opalescens* care, as well as Dr. Freeland Dunker, Kylie Lev, and other staff at the California Academy of Sciences for donating moon jelly polyps to the project. This work is in memory of our outstanding colleague Dr. Maryna P. Lesoway, who provided significant intellectual and emotional support in executing this work.

## Supplementary Figure Legends

**Figure S1.** Hierarchical clustering of Distal and Proximal Ceras tissue based on normalized expression profiles in DESeq2 in *Berghia stephanieae*.

**Figure S2.** PCA plot of Distal and Proximal Ceras tissue based on normalized expression profiles in DESeq2 in *Berghia stephanieae*.

**Figure S3.** Venn diagram of genes expressed in Distal and Proximal Ceras tissues in *Berghia stephanieae*. A total of 3,462 genes were not expressed in either tissue.

**Figure S4.** Hierarchical clustering of Distal (“cni”) and Proximal (“sub”) Ceras tissue based on normalized expression profiles in DESeq2 in *Hermissenda opalescens*.

**Figure S5.** PCA plot of Distal (“cni”) and Proximal (“sub”) Ceras tissue based on normalized expression profiles in DESeq2 in *Hermissenda opalescens*.

**Figure S6.** Venn diagram of genes expressed in Distal (“cni”) and Proximal (“sub”) Ceras tissues in *Hermissenda opalescens*. A total of 2,277 genes were not expressed in either tissue.

**Figure S7.** Hybridization chain reaction (HCR) controls. Early juveniles (A-A’), later juvenile (B-B’’), and adult (C-C’’) samples were incubated with only the amplifiers indicated (B1647and B2546) and no probes. Abbreviated: cb, cerata buds; cs, cnidosac; cp, cnidophage; dg, digestive gland.

**Figure S8.** Violin plots comparing VST transformed counts of genes in our analysis assigned particular GO terms. (A,D) GO:0007586, Digestion (119 genes total; 9 DE genes); (B,E) GO:0006897, Endocytosis (525 genes; 33 DE genes); (C,F) GO:0006955, Immune Response (838 genes; 52 DE genes). Plots are for subsets of genes associated with each of these three GO terms: (A-C) for all genes associated with each GO term, and (D-F) for only differentially expressed (DE) genes across both tissues associated with each GO term. Asterisks indicate level of significance between the two tissues (ns, p > 0.05; *, p <= 0.05; **, p <= 0.01; ***, p <= 0.001; ****, p <= 0.0001).

## Supplementary Table Legends

**Table S1.** Read mapping statistics for *Berghia stephanieae* and *Hermissenda opalescens*.

**Table S2.** All genes expressed in *Berghia stephanieae* samples that were found to be associated with the GO terms Endocytosis (GO:0006897) and Phagocytosis (GO:0006909) or child GO terms associated with these processes.

**Table S3.** Genes upregulated in the Distal Ceras in *Berghia stephanieae*.

**Table S4.** GO terms associated with genes upregulated in the Distal Ceras in *Berghia stephanieae*.

**Table S5.** Genes upregulated in the Proximal Ceras in *Berghia stephanieae*.

**Table S6.** GO terms associated with genes upregulated in the Proximal Ceras in *Berghia stephanieae*.

**Table S7.** GO terms used to generate violin plots in Figure 6.

**Table S8.** List of orthogroups containing genes determined to be upregulated in both *Berghia stephanieae* and *Hermissenda opalescens*. Annotations for the *B. stephanieae* genes within each orthogroup are also provided.

**Table S9.** Summary of HCR probes used in the manuscript, including the sequence used to generate each probe.

**Table S10.** Details for probes generated to detect *bsbhmt* expression.

**Table S11.** Details for probes generated to detect *bsferretin* expression.

**Table S12.** Details for probes generated to detect *bscol8a1* expression.

**Table S13.** Details for probes generated to detect *bsclect* expression.

## Notes

https://doi.org/10.5061/dryad.8pk0p2p00

https://github.com/goodgodric28/sequestration_genes/tree/main/sequestration_genes

### Competing Interest Statement

The authors have declared no competing interest.

