## Supplementary material for "A subset of conserved phagocytic genes are likely used for the intracellular theft of cnidarian stinging organelles in nudibranch gastropods": Figures S1-S8

Supplementary Figures S1-S3

### Figures S1-S3: Differential expression results

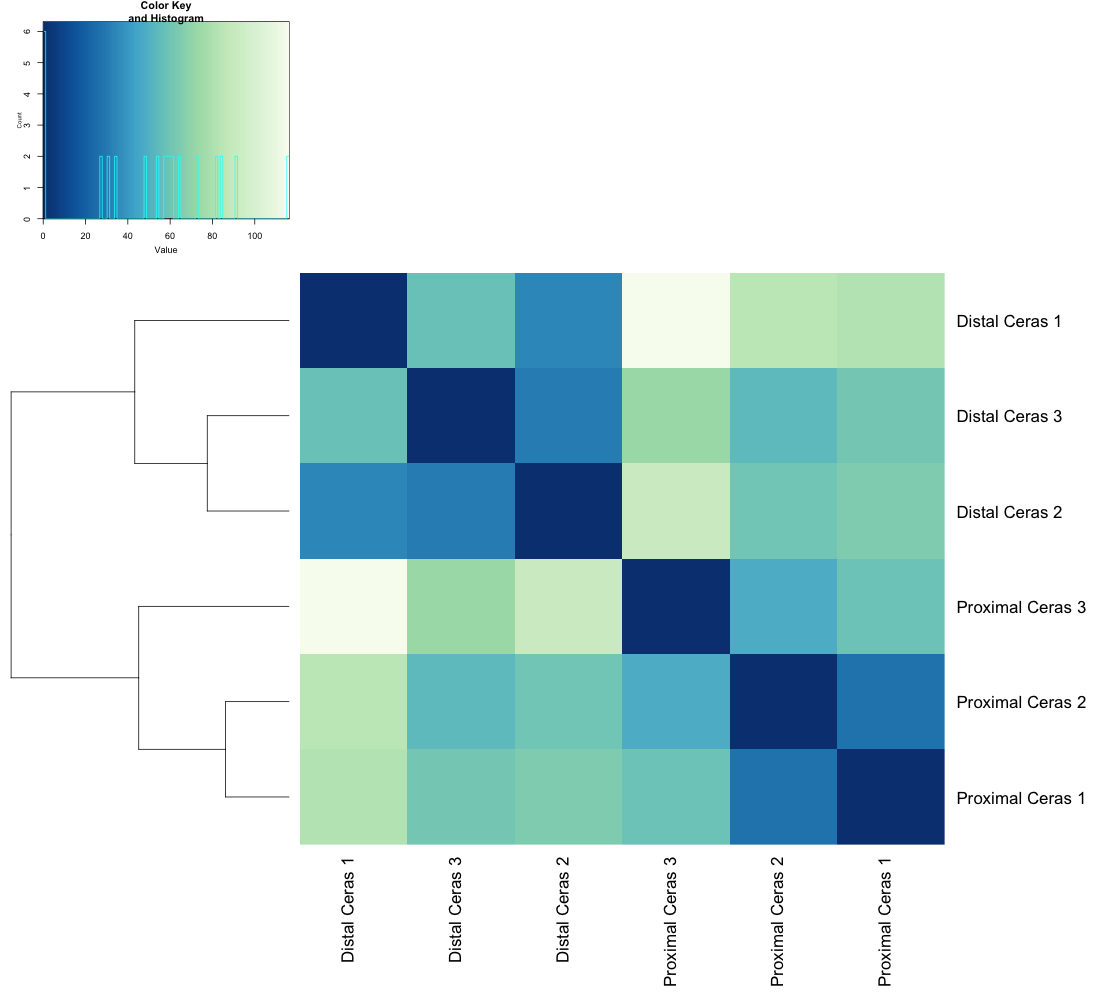

**Figure S1. Hierarchical clustering of Distal and Proximal Ceras tissue based on normalized expression profiles in DESeq2 in *Berghia stephanieae*.**

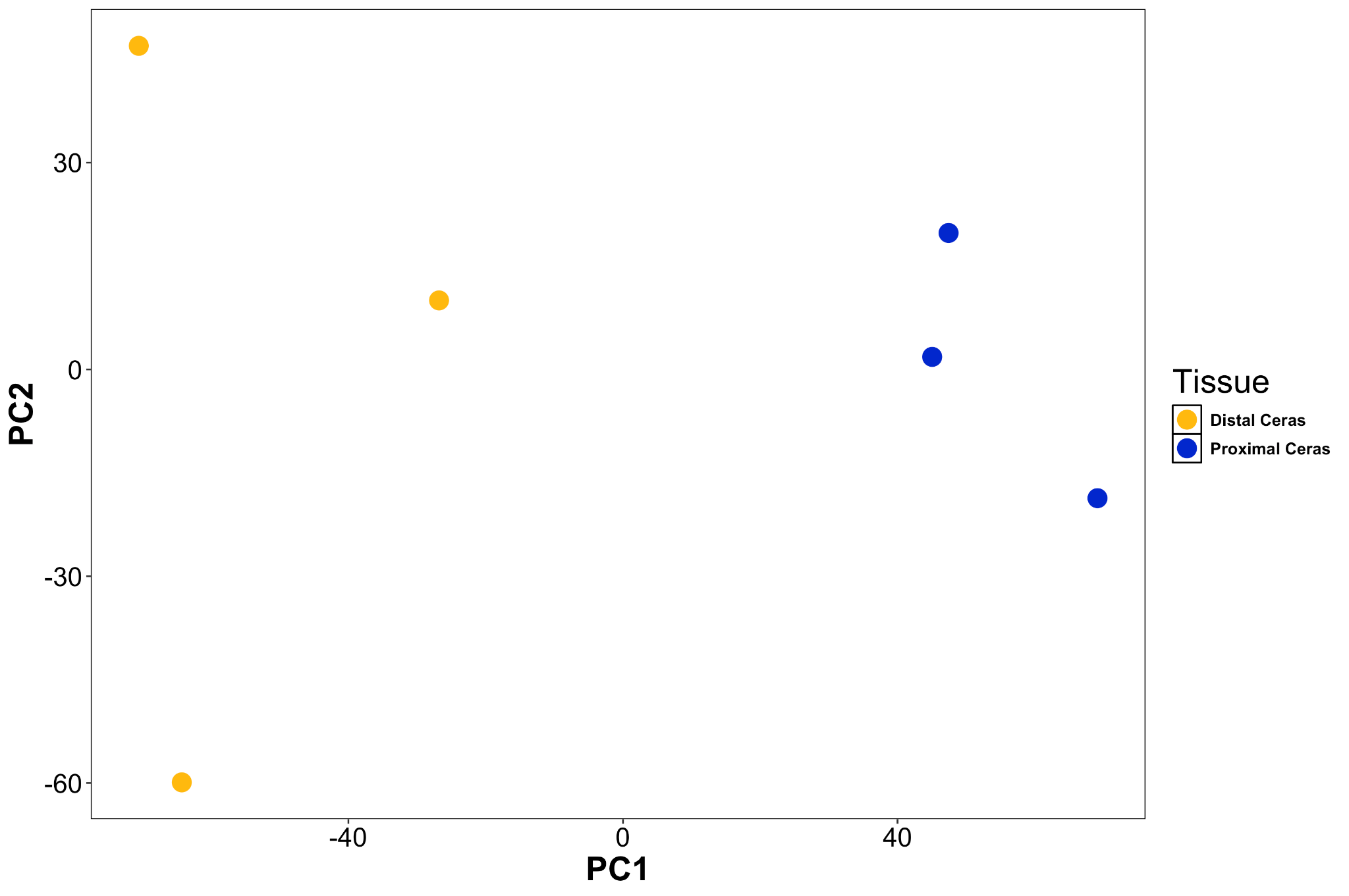

**Figure S2. PCA plot of Distal and Proximal Ceras tissue based on normalized expression profiles in DESeq2 in *Berghia stephanieae*.**

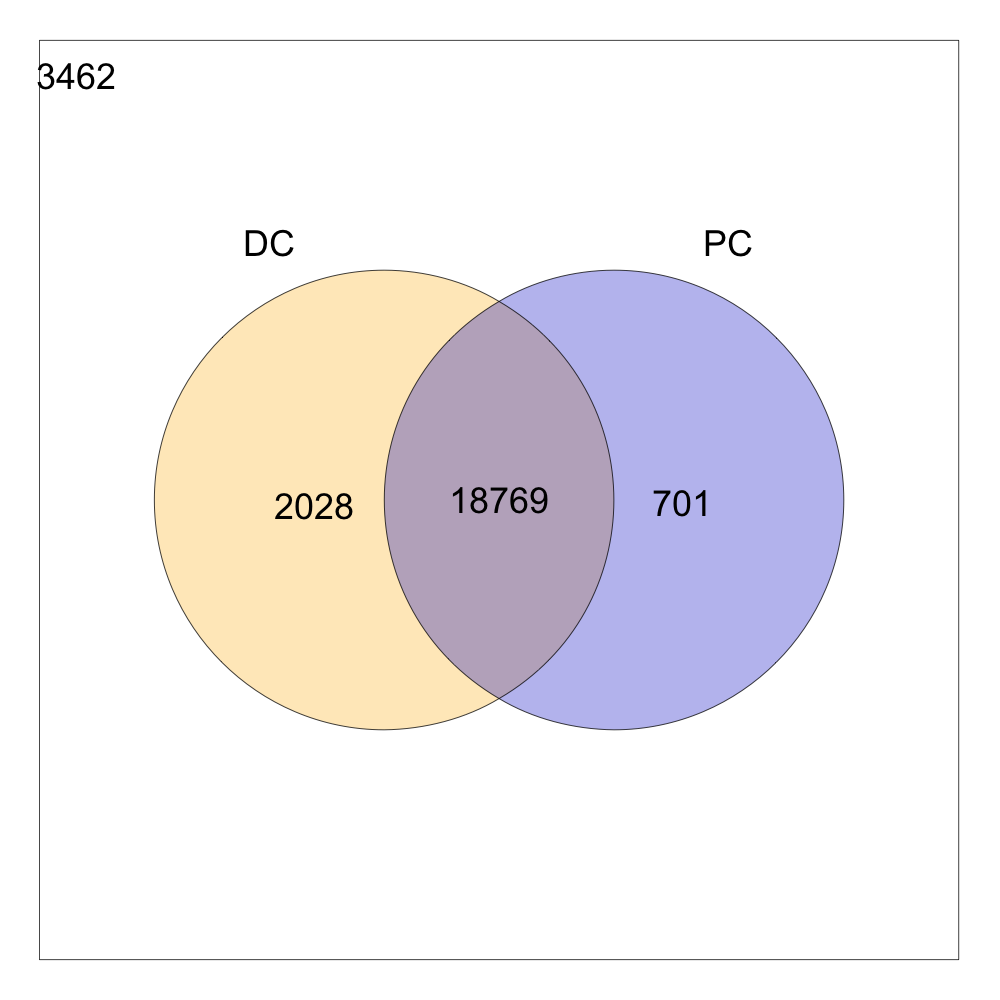

**Figure S3. Venn diagram of genes expressed in Distal and Proximal Ceras tissues in *Berghia stephanieae*.** A total of 3,462 genes were not expressed in either tissue.

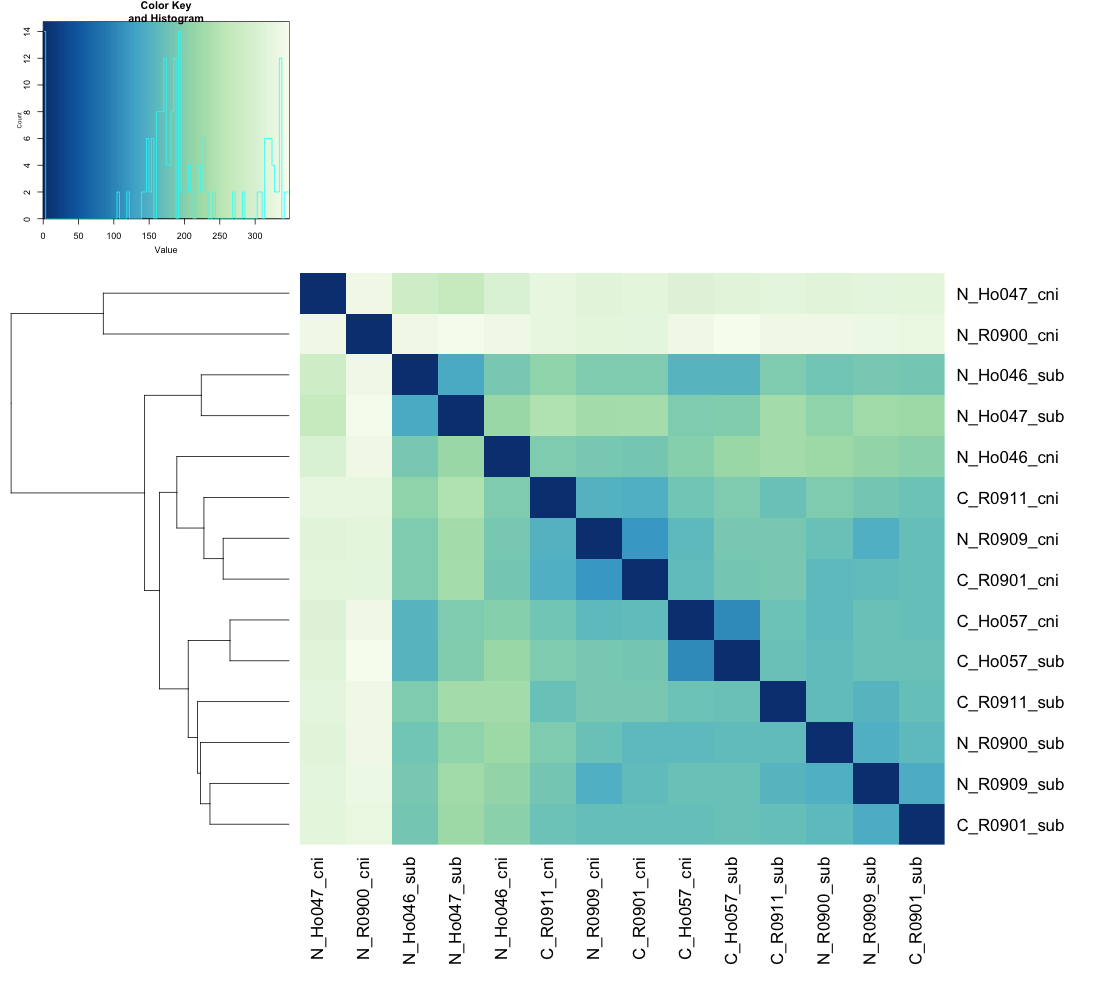

**Figure S4. Hierarchical clustering of Distal (“cni”) and Proximal (“sub”) Ceras tissue based on normalized expression profiles in DESeq2 in *Hermissenda opalescens*.**

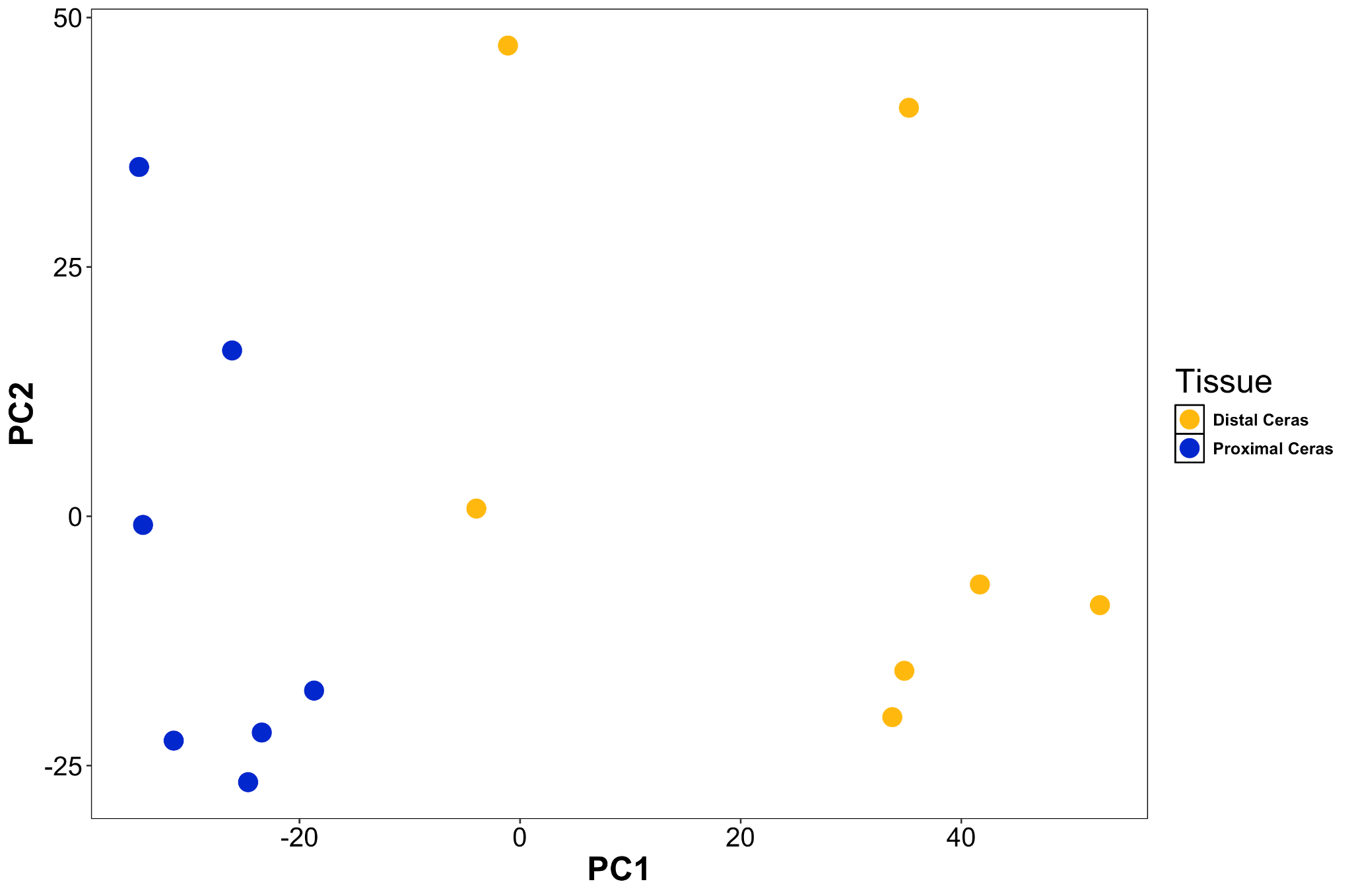

**Figure S5. PCA plot of Distal (“cni”) and Proximal (“sub”) Ceras tissue based on normalized expression profiles in DESeq2 in *Hermissenda opalescens*.**

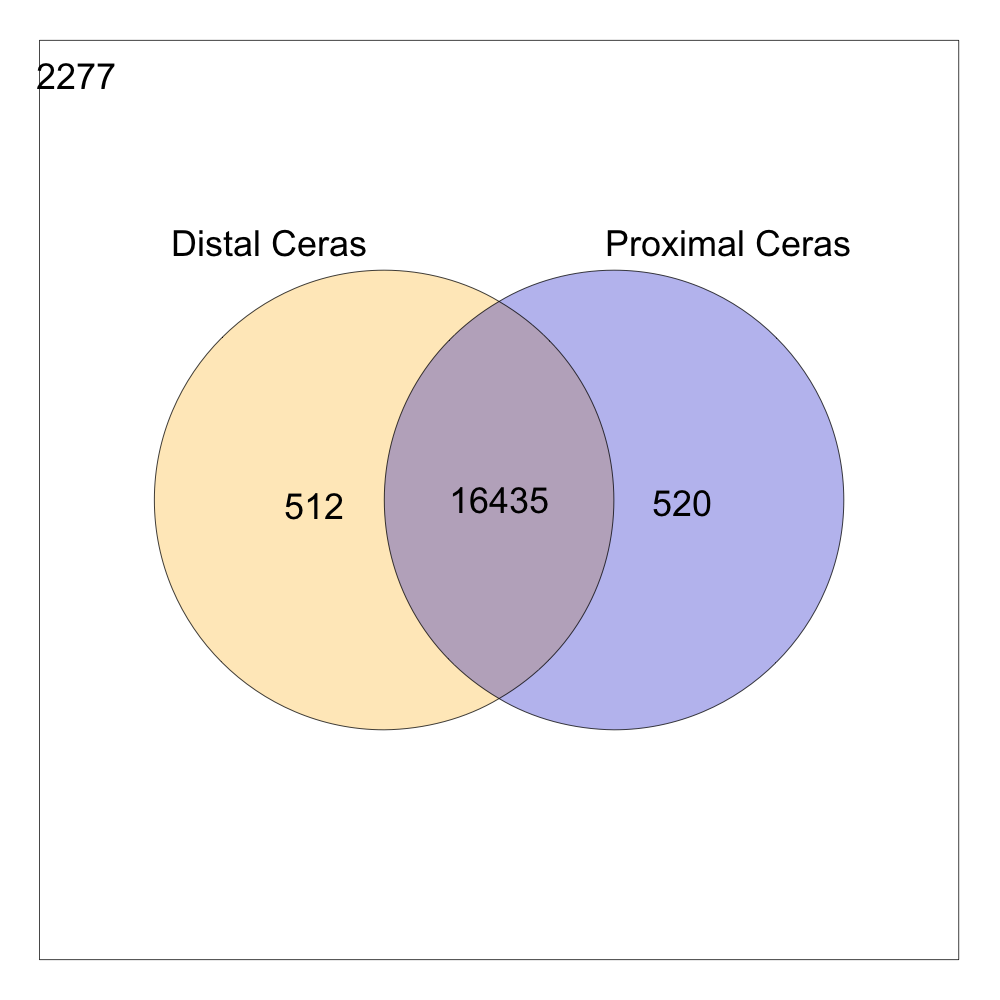

**Figure S6. Venn diagram of genes expressed in Distal (“cni”) and Proximal (“sub”) Ceras tissues in *Hermissenda opalescens*.** A total of 2,277 genes were not expressed in either tissue.

### Figure S7: HCR Controls

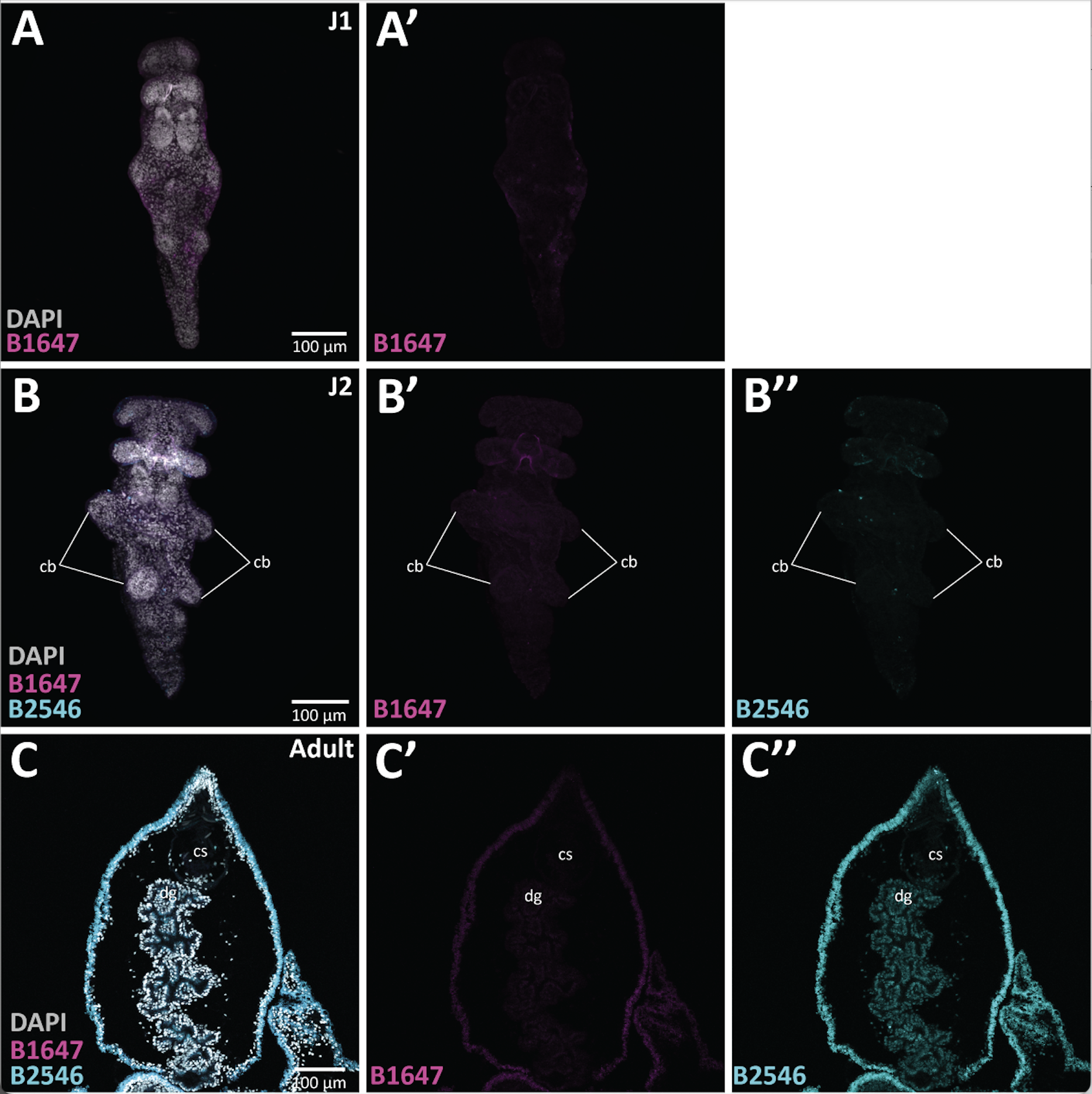

**Figure S7. Hybridization chain reaction (HCR) controls.** Early juveniles (A-A’), later juvenile (B-B’’), and adult (C-C’’) samples were incubated with only the amplifiers indicated (B1647and B2546) and no probes. Abbreviated: cb, cerata buds; cs, cnidosac; cp, cnidophage; dg, digestive gland.

### Figure S8: GO term comparisons

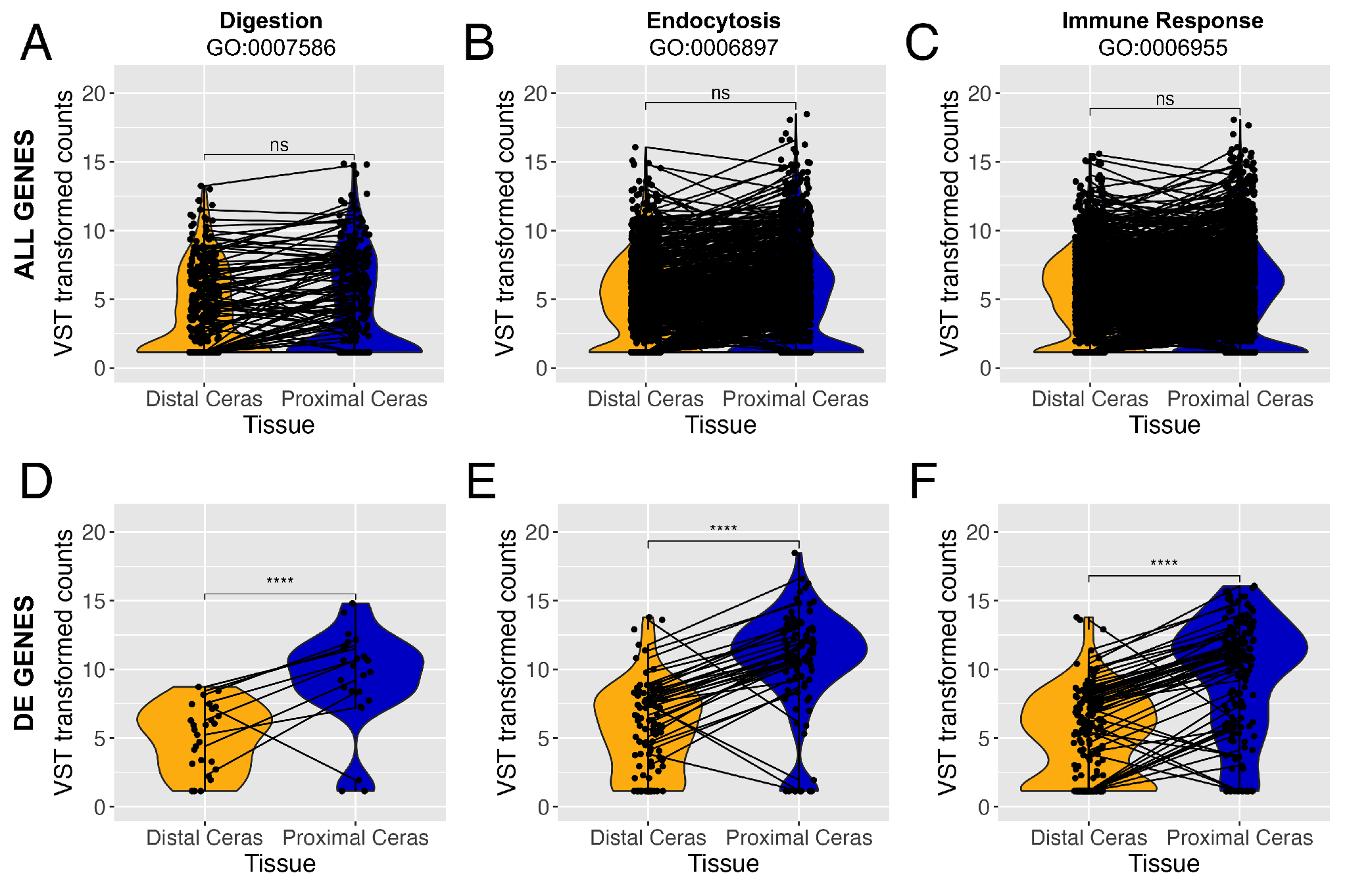

**Figure S8. Violin plots comparing VST transformed counts of genes in our analysis assigned particular GO terms.** (A,D) GO:0007586, Digestion (119 genes total; 9 DE genes); (B,E) GO:0006897, Endocytosis (525 genes; 33 DE genes); (C,F) GO:0006955, Immune Response (838 genes; 52 DE genes). Plots are for subsets of genes associated with each of these three GO terms: (A-C) for all genes associated with each GO term, and (D-F) for only differentially expressed (DE) genes across both tissues associated with each GO term. Asterisks indicate level of significance between the two tissues (ns, p > 0.05; *, p <= 0.05; **, p <= 0.01; ***, p <= 0.001; ****, p <= 0.0001).
